# Latent *Mycobacterium tuberculosis* infection provides protection for the host by changing the activation state of the innate immune system

**DOI:** 10.1101/561126

**Authors:** Johannes Nemeth, Gregory S. Olson, Alissa Rothchild, Ana Jahn, Dat Mai, Fergal Duffy, Jared Delahaye, Sanjay Srivatsan, Courtney Plumlee, Kevin Urdahl, Elizabeth Gold, Alan Diercks, Alan Aderem

## Abstract

An efficacious vaccine against adult tuberculosis (TB) remains elusive. Progress is hampered by an incomplete understanding of the immune mechanisms that protect against infection with *Mycobacterium tuberculosis* (Mtb), the causative agent of TB^1^. Over 90% of people who become infected with Mtb mount an immune response that contains the bacteria indefinitely, leading to a state known as “latent TB infection” (LTBI)^2^. A significant body of epidemiologic evidence indicates that LTBI protects against active TB after re-exposure, offering an intriguing avenue to identifying protective mechanisms^3, 4^. We show that in a mouse model, LTBI is highly protective against infection with Mtb for up to 100 days following aerosol challenge. LTBI mice are also protected against heterologous bacterial challenge (*Listeria monocytogenes*) and disseminated melanoma suggesting that protection is in part mediated by alterations in the activation state of the innate immune system. Protection is associated with elevated activation of alveolar macrophages (AM), the first cells that respond to inhaled Mtb, and accelerated recruitment of Mtb-specific T cells to the lung parenchyma upon aerosol challenge. Systems approaches, including transcriptome analysis of both naïve and infected AMs, as well as *ex vivo* functional assays, demonstrate that LTBI reconfigures the response of tissue resident AMs.. Furthermore, we demonstrate that both LTBI mice and latently infected humans show similar alterations in the relative proportions of circulating innate immune cells, suggesting that the same cellular changes observed in the LTBI mouse model are also occurring in humans. Therefore, we argue that under certain circumstances, LTBI could be beneficial to the host by providing protection against subsequent Mtb exposure.

## Introduction

The failure to identify mechanisms or correlates of protection against TB has hampered the development of vaccines or host-directed therapies to combat the world-wide disease burden which is estimated to be 10.4 million cases and 1.8 million deaths annually^5^. The immune mechanisms that protect against TB are evidently quite powerful: Due to the extraordinarily high prevalence of *Mycobacterium tuberculosis* (Mtb) (some estimates suggest that at least 25% of the world’s population has been exposed^2^) TB ranks as the deadliest infectious disease world-wide. However, the vast majority (∼90%) of individuals with an intact immune system are able to contain and control the infection for their lifetimes with no clinical symptoms leading to a state known as “latent TB infection” ^2^.

LTBI is clinically defined as the presence of an Mtb-specific adaptive immune response in the absence of signs and symptoms of disease. It is widely believed that the clinical definition encompasses a variety of disease states, often referred to as the “spectrum of TB disease”, including bacterial clearance, persistence of live bacteria contained by the immune system, and sub-clinical disease^6, 7^. It is clear that humans may harbor viable Mtb in a variety of tissues for decades without developing symptoms^8–11^. In both humans^12^ and non-human primates^13^, the lymphatic system plays key roles in the spectrum of disease by providing a reservoir for latent Mtb^8, 14^ and coordinating subsequent immune responses.

Both historical cohort studies and contemporary epidemiological studies demonstrate that LTBI is protective against re-infection^3, 15^. The phenomenon of a low-grade infection protecting against subsequent infections with the same pathogen has led to the development of almost all live vaccines currently in use, including those based on bacteria (BCG) or experimental vaccines against parasites (Leishmania)^16^. Likewise, continuous exposure to *Plasmodium falciparum*, the causative agent of malaria, and maintenance of low-grade parasitemia provides protection against high-density parasitemia and death in adults^17^. This phenomenon is often referred to as “concomitant immunity”^18^ or, in the case of malaria, “premunition”^17^. Despite the strong evidence that LTBI confers protection against reinfection, the underlying mechanisms have not been defined, in part due to the lack of a suitable animal model.

Recent studies in mice have shown that intradermally injected Mtb are asymptomatically contained in the draining lymph node for an extended period of time and have used this approach to model LTBI in humans with the goal of understanding failure of containment arising from immune suppression^19, 20^. Because the LTBI mouse model shares many features of latent TB in humans, we hypothesize that it reflects at least a portion of the disease spectrum. Here, we demonstrate that LTBI in mice confers significant protection against subsequent aerosol challenge, in part, by altering the innate immune response to Mtb.

## Results

### Intradermal Mtb-infection of mice recapitulates key aspects of human LTBI and is contained for up to 1 year

To establish the model of LTBI, we infected mice intradermally in the ear with 10,000 CFU of a commonly used virulent strain of Mtb, H37Rv^19^. Within 5 days the bacteria trafficked to the ipsilateral superficial cervical lymph nodes which, when extracted, were visibly enlarged. The bacterial burden in the draining lymph node was relatively stable for at least one year (Figure 1A) with minimal dissemination to the spleen and no dissemination to the lung (Figure S1). Approximately 75% of all spleens investigated were uninfected. If Mtb was detectable, the bacterial load was less than 1000 CFU per spleen (data not shown). We measured the circulating levels of 39 cytokines (including key inflammatory mediators such as IL-6 and TNF, (see Table S1) and found that none were significantly altered the first 6 weeks following the establishment of LTBI.

**Figure 1:**
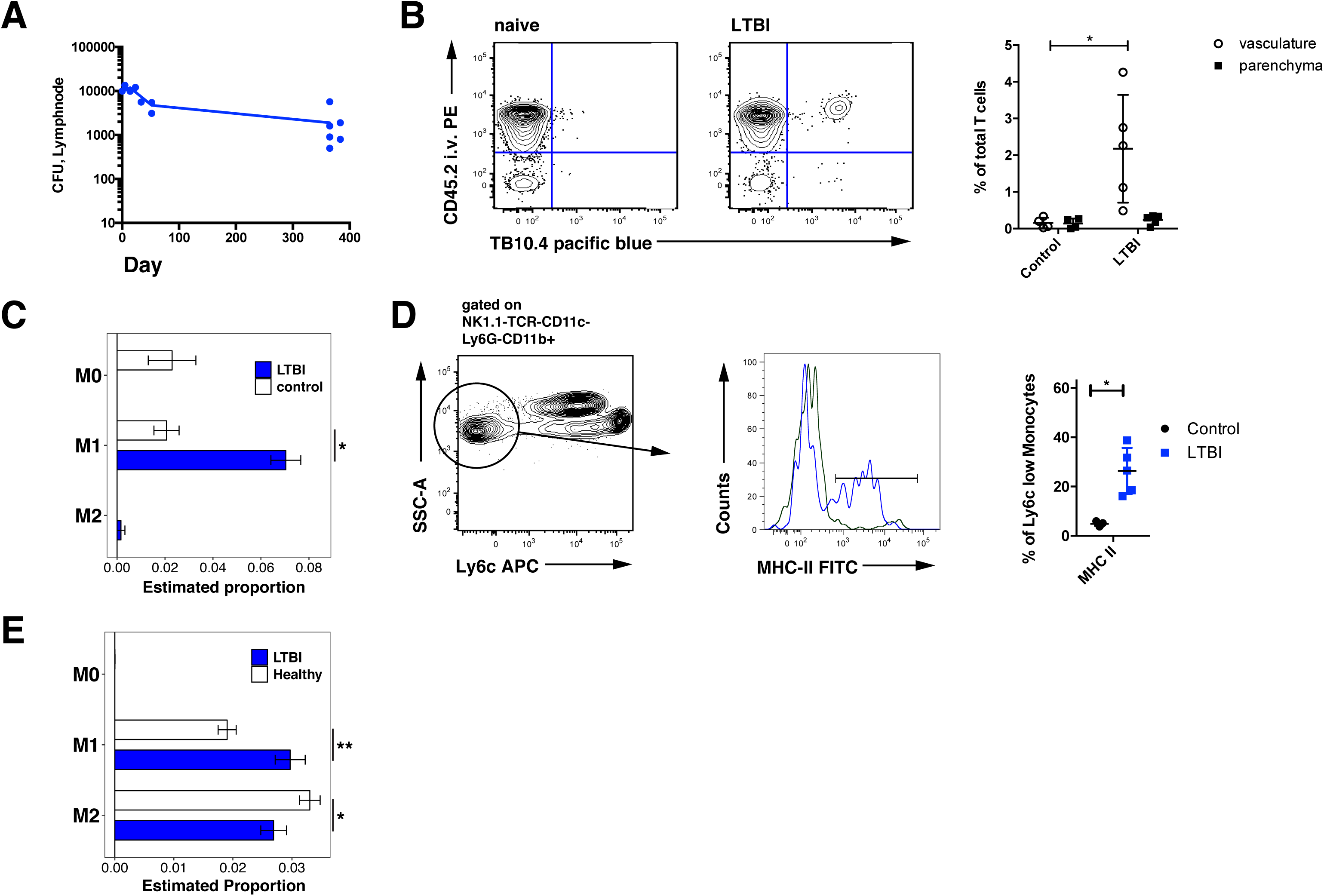
LTBI mice have an altered inflammatory milieu. **A:** Mice were inoculated intradermally in the ear and bacterial burden in the superficial cervical lymph nodes measured at the indicated time points by CFU assay. (1 representative experiment of 3 experiments, *n* = 3-6 mice/timepoint). **B:** LTBI was established as described in (**A**) and the relative proportions of CD3^+^CD8^+^TB10.4-tetramer-positive cells in the vasculature (CD45.2+) and parenchyma (CD45.2-) determined 42 days after inoculation by flow cytometry. Data are representative of 2 experiments with *n* = 4-5 mice/experiment. Statistical significance was determined by Student’s *t*-test. Error bars represent the mean with SD. **C:** LTBI was established as in (**A**) and whole-blood transcriptomes measured by RNA-seq in LTBI and control mice. The relative proportions of circulating M0, M1, and M2 macrophage type cells at day 42 following inoculation were determined by deconvolution analysis (See Methods) (4 mice/condition). Differences in proportions between control and LTBI mice with FDR < 0.05 are marked with “*”. **D:** LTBI was established as in (**A**) and expression of MHCII on Ly6c negative NK1.1^-^TCR^-^CD11c^-^Ly6G^-^CD11b^+^ monocytes in LTBI and control mice (5 mice/group), representative result of 2 independent experiments. **E:** Relative proportions of circulating M0, M1, and M2 macrophage type cells inferred by deconvolution analysis of whole-blood RNA-seq measurements on uninfected humans (n=117) or humans with LTBI (n=69). Error bars represent the SD. Significance was determined by *t*-Test with the Benjamini and Hochberg correction for multiple testing * FDR < 0.05, ** FDR < 0.001.

In humans, LTBI is clinically defined by the presence of an Mtb-specific T cell response in the absence of active disease. In LTBI mice, Mtb-specific T cells were detectable in the circulation from day 10 onward without recruitment to the lung parenchyma (Figure 1B). Furthermore, for at least one year, the mice displayed no overt systemic symptoms (e.g. weight loss or changes in coat or behavior) nor any local symptoms (including visible inflammation or irritation).

To assess more broadly whether LTBI changes the relative proportions of circulating immune cells, we performed whole blood RNA sequencing on LTBI mice and controls and applied a previously published transcriptome deconvolution approach based on a published deconvolution matrix^21, 22^. The most pronounced differences were increased frequencies of M1 macrophage type cells and monocytes in LTBI mice (Figure 1C). Flow-cytometry analysis of peripheral blood showed subtly but significantly elevated expression of MHC II on CD11b^+^Ly6C^low^ cells – the “tissue resident macrophages” in the blood compartment^23^ – from LTBI mice compared to controls (Figure 1D). To assess whether LTBI in humans is associated with changes in circulatory cellularity, we applied the same deconvolution approach to previously published whole blood transcriptome data from uninfected and latently infected humans^24^. In concordance with the mouse data, individuals with LTBI (*n* = 69) had on average a greater proportion M1 polarized macrophage type cells in the circulation than uninfected individuals (*n* = 117) (Figure 1E, S2). Across all samples, the estimated frequencies of M1 and M2 macrophage type cells in humans were inversely correlated with each other (Figure S3).

Taken together, these data indicate that intradermal Mtb-infection of mice recapitulates many key aspects of LTBI in humans, including circulating Mtb-specific T cells in the absence of disease symptoms or elevated levels of pro-inflammatory mediators in peripheral blood. Furthermore, LTBI in mice induces a skewing towards more M1 macrophages in the circulation that we also observed (and to our knowledge has not been previously described) in latently infected humans.

### Latently infected mice are strongly protected against aerosol challenge with Mtb and heterologous challenges

To test the hypothesis that LTBI is protective against aerosol infection, we infected mice with 50-100 CFU of Mtb H37Rv 8-10 weeks after the establishment of LTBI and measured bacterial burden in the lung and spleen at 14, 42, and 100 days after challenge. At each of these time points, the bacterial burden in both tissues was significantly lower in LTBI mice compared to controls (Figure 2A). We obtained similar results when mice were challenged 10 to 14 days after establishment of LTBI (Figure S4): LTBI mice had on average 18.4-fold (CI: 10.6-26.3) fewer bacteria in the lung as compared to controls measured 6 weeks after aerosol challenge across 6 independent experiments with a total of 60 mice. This level of protection exceeds that of intradermal inoculation with BCG which induces 10-fold protection at day 42 that wanes by day 100 and has minimal impact on bacterial burden within the first two weeks (Figure S5).

**Figure 2:**
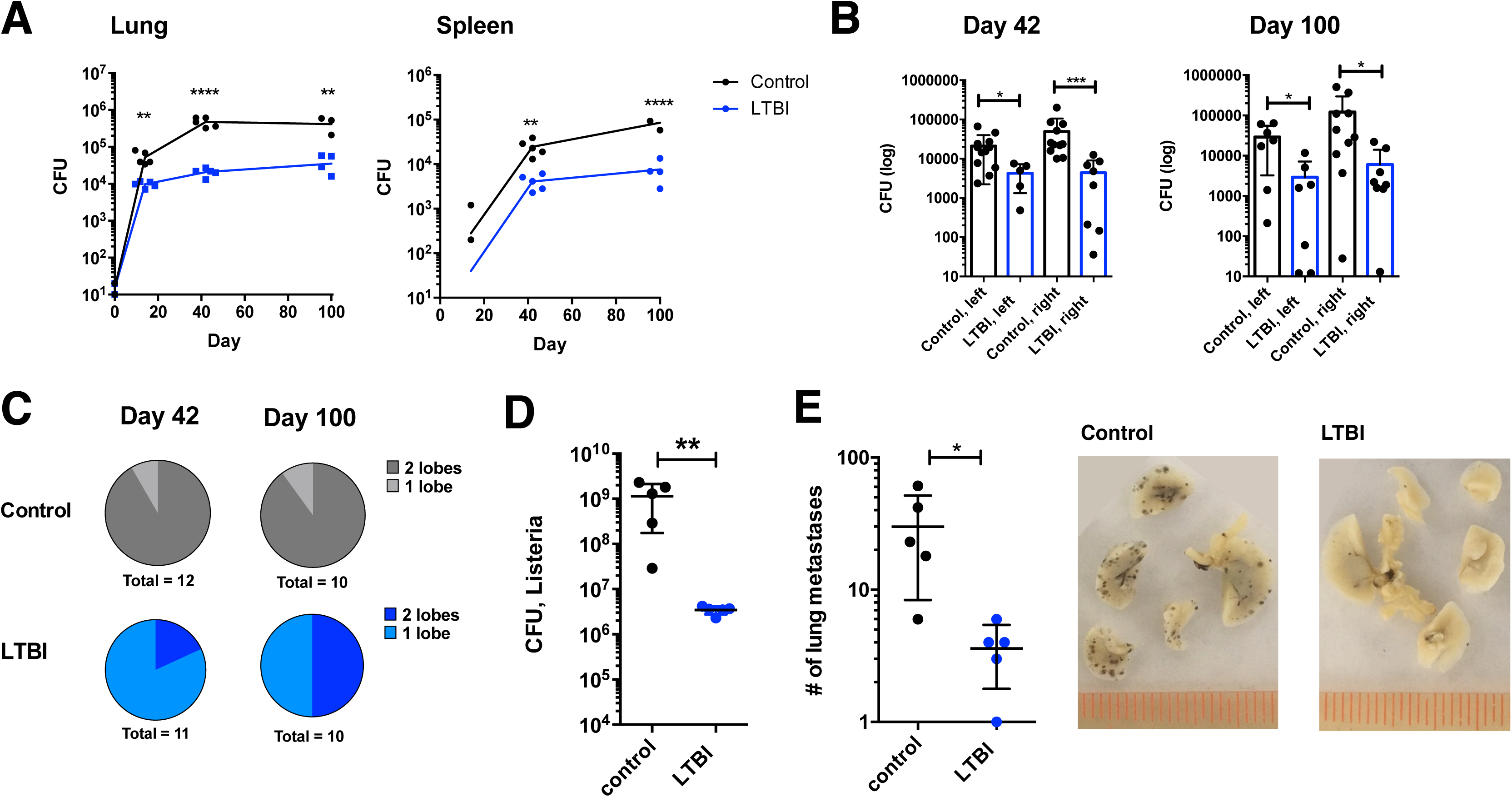
Latent TB infection provides sustained protection against aerosol challenge and heterologous challenges. **A:** 8 weeks after the establishment of LTBI, latent and control mice were challenged with 50-100 CFU of Kanamycin-resistant H37Rv via aerosol. Bacterial burdens in the lungs and spleen were measured by CFU at days 14, 42 and 100 following aerosol infection using plates containing Kanamycin. Plot shows representative data from one of two independent experiments (n = 4-5 mice/group/time-point). Significance was assessed by Student’s *t*-Test. Error bars represent the mean with SD; \*\**p* < 0.01, **** *p* < 0.0001. **B:** Naïve and LTBI mice (20/group) were infected with an average of 1 CFU of Mtb H37Rv via aerosol. Bacterial burdens on the left and right lobes of the lung were measured by CFU. Representative data from one of two independent experiments are presented. Uninfected mice were omitted in this graph. Statistical significance was determined using the Mann-Whitney test. Error bars represent the mean with SD. * *p* < 0.05, *** *p* < 0.001. **C:** Fractions of naïve and LTBI mice with detectable bacteria in one or two lobes of the lung in the experiment described in B. **D**: LTBI and control mice were challenged i.v. with 10^5^ CFU of *Listeria monocytogenes* and bacterial burden in the spleen measured 48 hours following infection by CFU. Representative data from one of two independent experiments with 4-5 mice/condition. **E:** LTBI and control mice were challenged i.v. with 1×10^6^ B16-F10 melanoma cells. Disease burden was quantified by counting the number of metastases (right panel: black spots) visible in the lung 10 days following challenge. Representative data of three replicate experiments with 4-5 mice/condition/time point. Significance was assessed by Student’s *t*-Test. Error bars represent the mean with SD; \**p* < 0.05.

Several lines of evidence suggest that human TB typically arises from infection with as few as 1-3 bacteria^11, 12^. We therefore examined the protective effect of LTBI against an “ultra-low dose” (ULD) infection of 1-3 CFU per mouse (see ^25^). At this dose, mice exhibit a wide range of outcomes, especially at later times, with bacterial burdens in the lung ranging over 4 orders of magnitude (Figure 2B). Furthermore, because most mice are infected with a single bacterium, the degree of dissemination can be inferred by measuring spread to the contralateral lobe. Consistent with the 50-100 CFU infections, LTBI mice challenged with 1-3 CFU showed substantially lower bacterial burdens and reduced dissemination in the lung compared to controls for more than 100 days following aerosol challenge (Figure 2C). These data demonstrate that LTBI mice are strongly protected against aerosol challenge with Mtb from as early as day 10 until at least 100 days following infection. To our knowledge, the early and sustained protective efficacy of LTBI is among the most profound that has been documented in the C57BL/6 mouse model.

Both adaptive and innate immune responses are required for control of Mtb. We reasoned that if the protective phenotype of the LTBI mice is fully explained by the presence of Mtb specific T cells (Figure 1B), these mice should be equally susceptible to other immune challenges that do not share specific antigens with Mtb. To test this, we challenged LTBI and control mice intravenously with 10^5^ CFU of *Listeria monocytogenes*, another intracellular bacterial pathogen. LTBI mice were significantly protected, displaying a greater than 10-fold reduction in bacterial burden in the spleen 48 hours following infection (Figure 2D). To confirm and extend these results, we inoculated LTBI and control mice with a non-bacterial immune challenge: a model of metastatic melanoma. Ten days following i.v. injection with melanoma cells, LTBI mice had approximately 10-fold fewer melanoma metastases in the lungs than controls (Figure 2E).

### The protective phenotype of latent mice is associated with modest tissue inflammation at baseline and an accelerated immune response

These data suggest that the protective phenotype induced by LTBI is mediated by immune mechanisms beyond Mtb-specific T cells. We hypothesized that LTBI modulates the innate immune system at baseline and in response to subsequent challenge leading to more efficient control of bacterial growth.

#### Prior to aerosol infection

Although LTBI did not induce elevated levels of cytokines in peripheral blood, we hypothesized that it might drive low-level, chronic immune activation in specific tissues. Therefore, we measured cytokine levels in the lungs and spleens of LTBI and control mice prior to aerosol challenge. During the first 6 weeks after establishment of LTBI, the levels of 7 cytokines and chemokines (of 39 assayed) were significantly elevated in the lungs of LTBI mice compared to controls and 5 were elevated in the spleen (Figure 3A). The most pronounced changes were increased levels of CCL3, CCL4, and CCL5 in the lung and IFNG and CCL4 in the spleen (Figure 3A). Relative levels of the detectable cytokines correlated well across tissues (supplemental Figure S6). We did not detect altered concentrations of classic inflammatory cytokines such as IL6 and TNF in any compartment tested. These results suggest that the localized, contained Mtb infection induces a mild and selective increase in local tissue inflammation.

**Figure 3:**
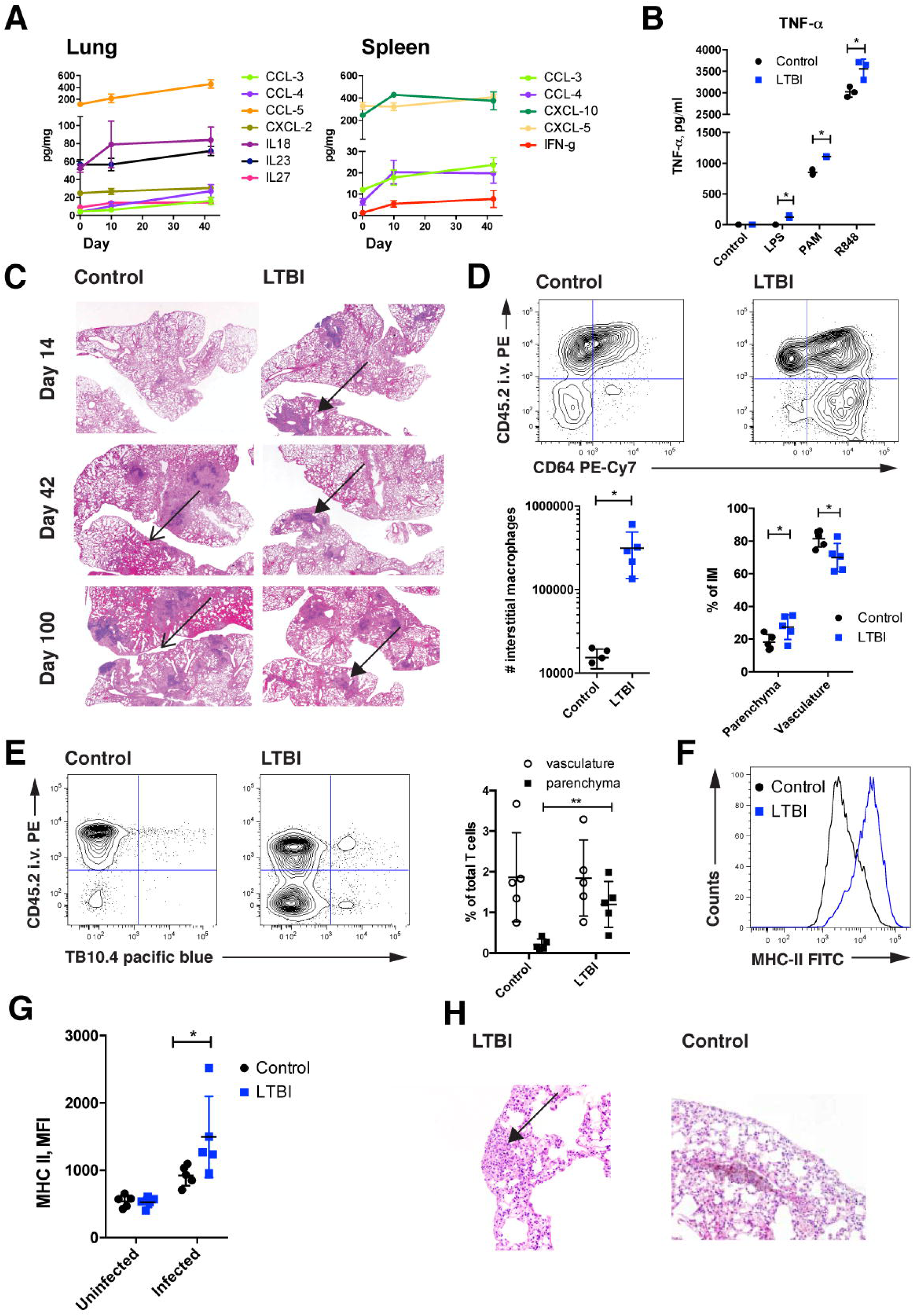
The protective phenotype in latent TB is associated with elevated tissue inflammation and an early immune response. **A:** Tissues were isolated from mice at 10 or 42 days following the establishment of LTBI or from naïve mice. Absolute levels of cytokines and chemokines significantly altered by LTBI during at least one time point in the lung or spleen (of 39 assayed) are plotted. Cytokine/chemokine amounts were normalized to total protein (*n* = 5 mice/time-point). A multiple *t*-test approach with FDR based correction was used to test for significance (p < 0.05). **B:** CD11b+ cells isolated from the lungs of LTBI or control mice were stimulated for 6 hours with the indicated TLR agonists and secreted TNF-α measured by ELISA (LPS 10 ng/mL, PAM3 300 ng/mL, R848 100 ng/mL). Representative data from 3 independent experiments with cells from 3-5 mice pooled per timepoint. Error bars represent the mean with SD. **C:** H&E stained lung sections from control and LTBI mice at the indicated time-points following aerosol challenge. Regions of infiltration by immune cells (filled arrow heads) and intrapulmonary hemorrhages (open arrow heads) are indicated. The corresponding quantitative pathology assessments are presented in Table S2. Representative images from 5 mice/group/time-point are displayed at 2x. **D:** Representative cytometry plot showing i.v. labeled, CD11b^+^CD11c^+^CD64^+^Siglec-F^-^ interstitial macrophages (IM) being recruited to the parenchyma. Latent mice show significantly higher absolute and relative numbers of IM at day 14 as compared to controls (*n* = 5, Error bars represent the mean with SD). Representative experiment from two independent experiments. **E:** Representative cytometry plot showing i.v. labeled CD3^+^CD8^+^TB10.4^+^ cells being recruited into the parenchyma. *t*-Tests was used for testing of significance. Error bars represent the mean with SD.* *p* < 0.05, ** *p* < 0.001. **F:** Expression of MHCII on alveolar macrophages (CD11b^int^CD11c^+^CD64^+^Siglec-F^+^) from control and LTBI mice isolated from the lung by BAL at 10 days following aerosol challenge. Data are representative of two independent experiments with 5 mice/condition. **G:** MHCII expression on infected splenic macrophages 2 hours after i.v. challenge with GFP-expressing *Listeria monocytogenes*. Representative of two independent experiments with 4-5 mice/condition. **H:** Representative H&E staining of a section from LTBI mice challenged with melanoma showing histiocytic accumulation (arrow heads).

#### After subsequent challenge

We first assessed the overall responsiveness of myeloid cell populations in the lung by isolating CD11b^+^ cells from LTBI and control mice and stimulating them for 6 hours with TLR-agonists. Myeloid cells from LTBI mice were more responsive to all stimuli tested as measured by TNF-α secretion (Figure 3B). The myeloid populations in the lung of LTBI and control mice were not significantly different (Figure S7).

Next, we investigated the immune response after aerosol Mtb infection by both histology of fixed lung sections and flow cytometry of single cells. In contrast to control mice, which had essentially no histologically detectable response to aerosol Mtb-challenge (100 CFU) at day 14 following infection, LTBI mice exhibited well-formed pulmonary lesions that remained stable for at least 100 days (Figure 3C). By day 42, control mice showed significantly more lesions and pulmonary hemorrhages than LTBI mice and this damage continued to progress through day 100 (Figure 3C, Table S2).

To identify the myeloid cells involved in the early response we observed histologically, we measured the recruitment of immune cells to the lung by flow cytometry at early timepoints (days 10-14) after aerosol infection. The magnitude of recruitment to the lung parenchyma of both interstitial macrophages and Mtb-specific T cells, was higher at this early timepoint in LTBI mice compared to controls (Figure 3D-E). In addition, alveolar macrophages (AMs) from LTBI mice were more highly activated as measured by MHCII expression following aerosol challenge (Figure 3F).

These results suggest that a key feature of LTBI is an accelerated, early immune response that reduces the time during which Mtb is able to replicate unchallenged. Furthermore, LTBI mice are able to maintain a protective response with minimal progression of immune pathology for a prolonged period of time.

#### Heterologous challenges

We hypothesized that heterologous challenges would also be associated with increased myeloid activation. To test this, we infected mice intravenously with 10^7^ GFP-expressing *Listeria monocytogenes* and measured the activation state of splenic macrophages after 2 hours. In concordance with our hypothesis, infected splenic macrophages from LTBI mice upregulated MHCII more strongly than macrophages from controls, suggesting that they were able to respond more robustly to the bacteria (Figure 3G). In addition, we evaluated tissue sections from the lung of LTBI or control mice challenged with i.v. melanoma. Strikingly, histiocytic and lymphocytic infiltrates were detectable in the lungs of 3 out of 5 LTBI mice and absent in control mice (Figure 3H, Table S3).

### Latent Mtb infection alters the response of alveolar macrophages to aerosol Mtb challenge

Since the protective phenotype in LTBI mice is strongly associated with early recruitment of inflammatory cells to the site of infection (Figure 3) and there is evidence of low-level tissue inflammation prior to challenge (Figure 3A), we hypothesized that LTBI affects the activation status and initial response of AMs (the lung resident macrophage and the first cells to be infected with Mtb^16^).

#### Prior to aerosol infection

While the total number of CD45^+^ cells and the fractions of both myeloid and T cells in the lung were unaffected by LTBI (Figure S7), we consistently measured elevated expression of MHCII and FcγRI on AMs (Figure 4A), suggesting that their activation state was altered by continuous exposure to low-level inflammation (Figure 3A). Expression of CD11c was unaffected by LTBI (Figure 4A). To investigate these cells at a more global level, we performed RNA sequencing on AMs isolated from LTBI and control mice. Interestingly, this RNA-seq analysis demonstrated that the transcriptomes of AMs from control and LTBI mice did not differ significantly prior to aerosol challenge (Figure 4B).

**Figure 4:**
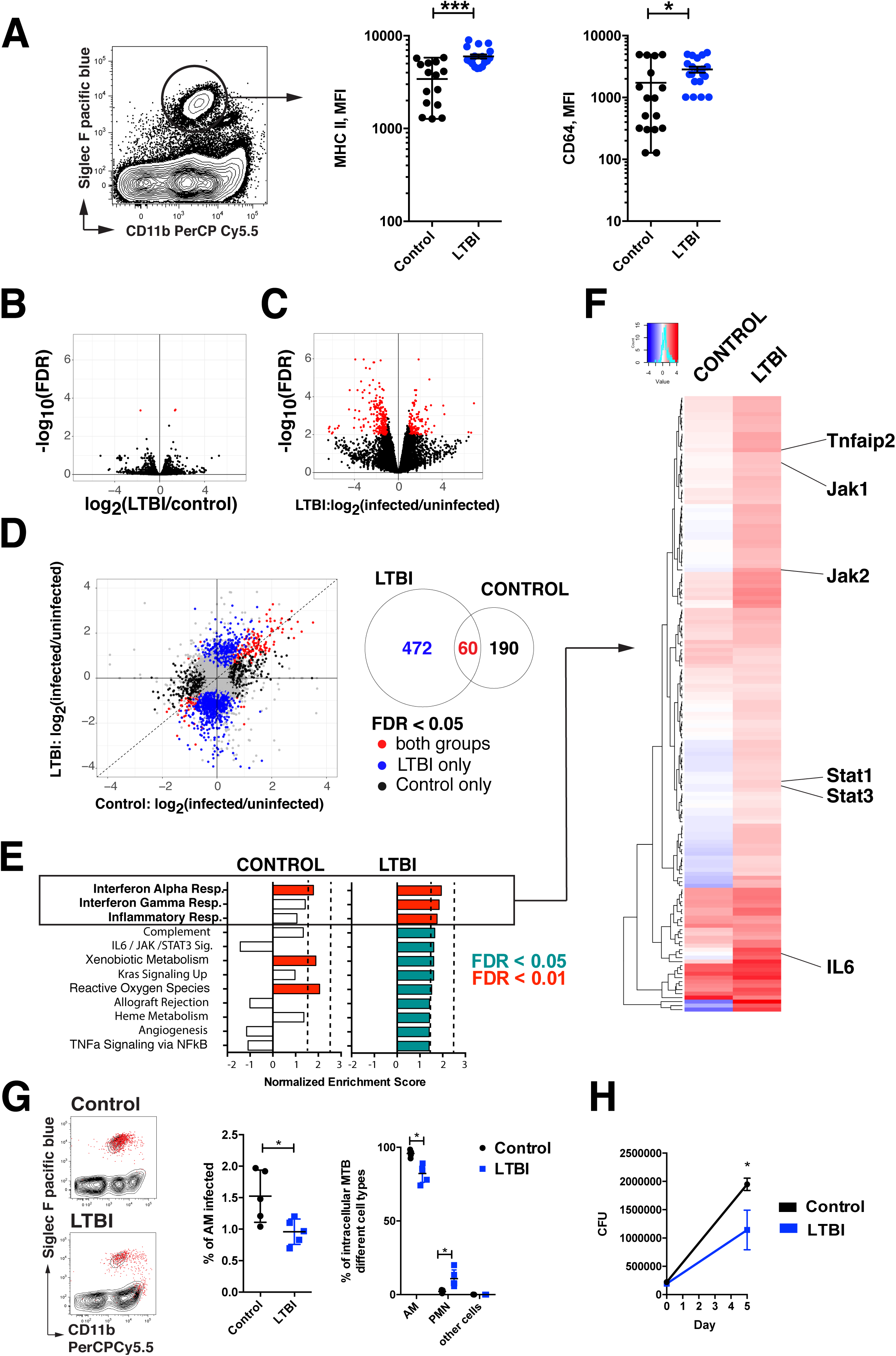
Alveolar macrophages are rewired by LTBI. **A:** Flow cytometry analysis of alveolar macrophages from control and LTBI mice. (Left panel) Live, CD11b^int^CD11c^+^CD64^+^Siglec-F^+^ were gated on Cd11b and Siglec F to define AMs. (Right panels) Expression of the indicated markers on AMs quantified by MFI. Significance was determined by Student’s *t*-Test; * p < 0.05. Error bars represent the mean and SEM. (Data pooled from 4 independent experiments with 3-5 mice/experiment). **B:** AMs from LTBI mice and controls were isolated by FACS and the transcriptome assessed by RNA sequencing. Red dots represent transcripts that were significantly differentially expressed between conditions (|log_2_(fold-change)| > 1 and FDR < 0.05). Each point represents the average of 3 biological replicates. **C:** Infected AMs from LTBI mice and controls were isolated by FACS 24 hours after aerosol challenge with ∼3000 CFU mEmerald-expressing Mtb. Red dots represent transcripts that were significantly differentially expressed between conditions (log2(fold-change) > 1 and FDR < 0.05). Each dot represents the average of two biological replicates, each of which consists of BAL pooled from 10 mice. **D:** (left panel) Transcripts differentially regulated (as defined in C) in AMs from naïve mice compared to AMs from LTBI mice. Transcripts responding to infection are color-coded by whether they are differentially expressed in: blue = LTBI only, black = control only, red = both conditions. (right panel) VENN diagram showing the number of differentially expressed transcripts in each group. **E:** Enrichment scores for the 10 most enriched pathways defined by GSEA of transcripts altered by Mtb infection at 24 hours in AMs from control (right panel) and LTBI (left panel) mice are shown. **F:** Heatmap showing the fold changes of unique, significantly enriched leading edge genes from the GSEA. Key transcripts are highlighted on the right. **G:** Control and LTBI mice were infected with ∼3000 CFU of mEmerald-expressing Mtb and the distribution of infected cell types determined by flow cytometry at 10 days following challenge. (left panel) Flow cytometry plot showing mEmerald infected cells (red) overlaid on plots gated on AM and PMNs. Quantification of (middle panel) the fraction of Mtb in various cell types and (right panel) the fraction of AMs infected. Representative data for one of three replicate experiments (n=4-5 mice/condition/experiment) are shown. Significance was determined by Student’s *t*-Test; * p < 0.05. Error bars represent the mean and SD. **H:** AMs from LTBI mice and controls were harvested by bronchioalveolar lavage and infected with H37Rv ex vivo at an MOI of 1 in triplicates. Bacterial burden was measured by CFU 5 days following infection. One out of two representative experiments, *n* = 4-5 mice/condition/time point. *t*-Tests was used for testing of significance. * p < 0.05. Error bars represent the mean with SD.

#### Following aerosol infection

To assess the immediate response of AMs to infection with Mtb, we challenged mice with a high dose (∼3000 CFU) of mEmerald-expressing bacteria. In concordance with previous studies, at 24 hours following infection the bacteria were predominantly contained within AMs^24, 25^. LTBI had no impact on the proportions of cell-types infected (Figure S8). In other studies (currently submitted), we have shown that MTB-infected AMs display a highly delayed pro-inflammatory reponse over the first 10 days of infection^26^. RNA-seq analysis of AMs isolated from control and latent animals 24 hours after aerosol challenge demonstrated a dramatically altered response to Mtb-infection in AMs from LTBI mice compared to controls (Figure 4C,D). Gene Set Enrichment Analysis (GSEA)^27^ of genes differentially expressed following infection showed that, in contrast to AMs from control mice, AMs from LTBI mice upregulated transcripts associated with inflammation, the Interferon-γ pathway and the IL-6 JAK-STAT pathways (Figure 4E,F).

The bacterial burden in LTBI mice at 10 days following high-dose (∼3000 CFU) aerosol infection, was approximately 3-fold lower than in control mice (Figure S9). The majority of bacteria were in AMs (Figure 4G) and significantly fewer AMs were infected in LTBI mice (Figure 4G). AMs isolated by BAL from LTBI mice were better able to control Mtb infection *ex vivo* compared to controls suggesting that the enhanced response of AMs from LTBI mice is at least in part cell-intrinsic. (Figure 4H).

Numerous recent studies have suggested that prior or ongoing infection can alter the capacity of innate immune cells to respond to subsequent encounters with pathogens, a phenomenon that has been termed “trained immunity” and has been shown in some cases to be reflected in epigenetic modifications^28^. Therefore, we used ATAC-seq to measure genome-wide changes in chromatin accessibility induced by LTBI. Surprisingly, we observed only modest changes in chromatin accessibility that did not correlate with differentially expressed genes, suggesting that this mechanism cannot account for the enhanced response of AMs to infection (Figure S10).

Taken together, these data suggest that changes in the local, inflammatory environment reprograms the transcriptional responses of AMs and allows for improved control of subsequent aerosol challenges.

### The protective phenotype of LTBI mice is diminished by antibiotic therapy

In the absence of long-lasting epigenetic changes, we hypothesized that the protective effect of LTBI results in part from the continuous exposure of innate cells to low-level cytokines secreted by immune cells containing the bacteria and thus would be diminished by antibiotic treatment. Therefore, we treated LTBI mice and controls for 6 weeks with Isoniazid and Rifampicin prior to aerosol challenge and assessed the impact on bacterial burden. Treatment efficacy was confirmed by culture of lymph node, spleen, and lung lysates from a dedicated set of mice (*n* = 2-3, two independent experiments, data not shown). Antibiotic treatment significantly diminished the ability of LTBI mice to contain bacterial growth in the lung over the first 6 weeks (Figure 5A). At 10 days following aerosol challenge, treated LTBI mice had lower numbers of Mtb-specific T cells in the circulation and in the lung parenchyma compared to untreated LTBI mice (Figure 5B). 5B In addition, treatment significantly reduced the expansion of Mtb-specific T cells in response to infection (Figure S11). These data suggest that the protective phenotype induced by LTBI is dependent on live bacteria.

**Figure 5:**
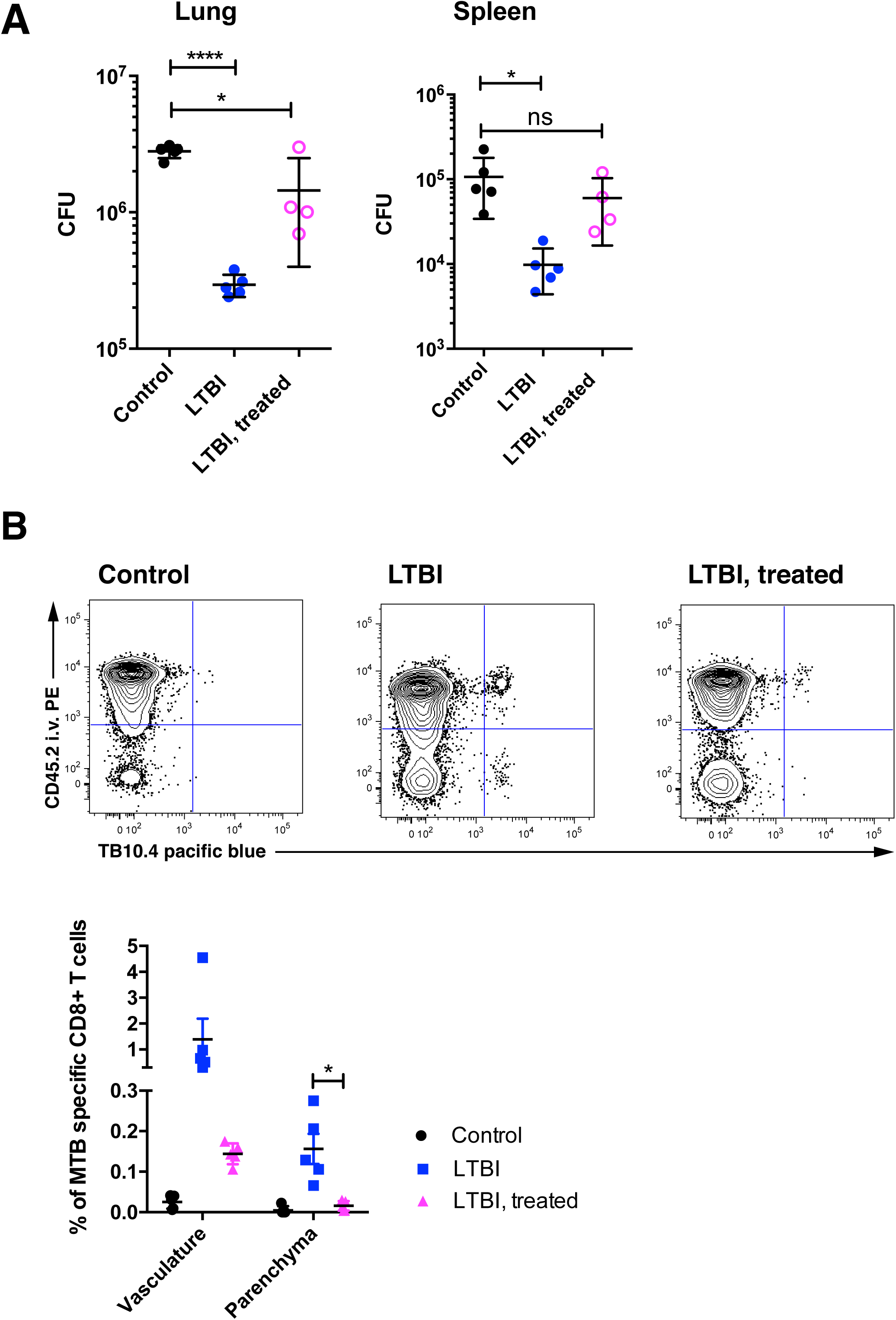
Treatment of latent infection reduces the protective effect of LTBI. **A:** Bacterial load in lungs and spleen 6 weeks after aerosol challenge with 100 CFU Mtb H37Rv for the indicated conditions. “LTBI, treated” and “Control” mice received INH/RIF for 6 weeks prior to aerosol challenge. Representative data from one of two independent experiments with 4-5 mice/condition. **B:** Representative flow cytometry plot showing recruitment of i.v. labeled CD3^+^CD8^+^TB10.4^+^ cells to the parenchyma in control mice, LTBI mice, and treated LTBI mice 10 days after aerosol challenge.

## Discussion

In this study, we have demonstrated that latent infection with virulent Mtb protects mice against subsequent aerosol challenge and that a large component of this protection arises from elevated activation of the innate immune system. Our data suggest that low-level inflammation within the lung enables resident AMs to respond more robustly to infection with Mtb. The increased innate immune response in LTBI mice also confers significant protection against heterologous challenges as diverse as *Listeria monocytogenes* and metastatic melanoma.

The mouse remains the premier small animal model for immunological research due to the extensive availability of molecular reagents, a large body of historical data, and relatively low cost. Much of our current understanding of TB immunology had its origins in the mouse model before subsequent validation in humans, e.g., the critical roles for CD4 T cells^29^, IFN_G_^30, 31^, IL-12^32^, and TNF^30^.

Furthermore, the mouse continues to play an essential role in the development of new TB vaccine strategies^33^. The protective effect of LTBI manifests as a reduced bacterial burden as early as 10 days following aerosol challenge that is maintained for at least 3 months after challenge. To our knowledge, this protection is as rapidly acting and durable as any that has been demonstrated in the mouse. Therefore, we argue that the LTBI mouse model offers an important opportunity to identify and dissect mechanisms of protection against TB.

Enhanced innate immune responses resulting from prior exposure to pathogens, a phenomenon termed “trained immunity”, is often reflected in epigenetic and transcriptional changes in macrophages and has been shown to be important for protection against Mtb in a model of high dose i.v. BCG vaccination^34^. The protective effect of i.v. BCG depends on the presence of bacteria in the bone marrow, affects the development of myeloid derived macrophages, and is transferrable via macrophages in the absence of live BCG^34^. In the LTBI model presented here, we detect profound changes in the responsiveness of tissue resident AMs, a long-lived cell type derived from fetal monocytes^35^. Furthermore, in the LTBI model, protection depends on live mycobacteria presumably via the continuous interaction between pathogen and host.

Tissue resident AMs from naïve mice have been reported to be niches for Mtb growth, in contrast to myeloid derived macrophages which are better able to control bacterial replication^36^. Mtb-infected AMs from LTBI mice upregulate many of the inflammatory and interferon pathways that are activated in myeloid-derived macrophages but not in Mtb-infected AMs from control mice (Figure 4E and ^19^). Thus, LTBI reprograms tissue resident AMs, the cells first infected by inhaled Mtb, to a more bactericidal phenotype that is typically associated with recruited macrophages.

Several mouse models demonstrate that infection or inoculation with live vaccines (including latent Herpes virus infection, BCG vaccination, and exposure to typical mouse pathogens) confers substantial protection against heterologous challenges by altering the activation state of the innate immune system^37–39^. Although the potential role in humans of LTBI-mediated heterologous protection has not been rigorously assessed, beneficial heterologous effects of BCG vaccination have been reported: overall mortality of newborns is reduced by more than would be expected from the protection afforded by BCG against TB alone^40^. Both local administration of BCG as well as BCG vaccination restrain melanoma development in humans^41, 42^. BCG-vaccination of mice has been reported to enhance the cytotoxic capacity of isolated macrophages against melanoma cells *in vitro*^43^. Furthermore, skewing of macrophages from M2 to M1 has been associated with control of melanoma *in vivo*^13, 14^.

Antibiotic treatment of LTBI mice reduced the numbers of Mtb-specific T cells in the circulation and in the lung parenchyma following aerosol challenge. Surprisingly, treatment also eliminated specific T cell expansion. We hypothesize that eliminating the ongoing contained infection also eliminated the elevated activation of the innate immune system resulting from LTBI and thereby reduced appropriate instruction of the adaptive response. However, more detailed studies are required to elucidate the contributions of memory T cells and reprogrammed macrophages to the protective effect of LTBI.

Preventative antibiotic treatment of patients diagnosed with latent TB is regarded as a cornerstone of TB control. However, preventive therapy is only effective in the first two years after infection and fails to reduce TB incidence if applied without risk-stratification^4, 44^. Counterintuitively, recent epidemiological data suggest that individuals who have been treated for active TB are more susceptible to re-infection and progress more rapidly following re-infection, suggesting that clearing latent infection with preventive therapy might be counterproductive in a high transmission setting^15, 45^. This phenomenon has been observed for malaria: treatment of low-grade *Plasmodium falciparum* parasitemia and subsequent loss of natural immunity can lead to hyper-mortality in epidemic regions^46^. In both cases, the protective mechanisms that are undermined by treatment are incompletely understood. A detailed examination of antibiotic treatment in the LTBI mouse model, in conjunction with human studies, may help to uncover these mechanisms and ultimately inform improved clinical treatment of TB in high-incidence settings.

## Methods

### Mice

C57BL/6 were purchased from the Jackson Laboratory. All mice were housed and bred under specific pathogen free conditions at the Center for Infectious Disease Research. All experimental protocols involving animals were approved by the Institutional Animal Care and Use Committee of the Center for Infectious Disease Research.

### Tissue Culture

Alveolar macrophages were cultured in complete RPMI [cRPMI; plus 10%(vol/vol) FBS, 2 mM l-glutamine, penicillin, and streptomycin] for 24 hours after isolation by bronchoalveolar lavage (BAL). In vitro infections were performed in cRPMI without antibiotics in biological triplicate (cells from independent mice). The bacterial load at day 5 was determined by plating serial dilutions of cells lysed in 1% Triton-X and diluted in 0.1% Tween-80 PBS.

### Establishment of LTBI via intradermal infection of the ear, aerosol Infections, and enumeration of bacterial load

Intradermal infections to establish were performed as described previously^19^ with the following modifications: 10,000 CFU of Mtb (H37Rv) in logarithmic phase growth in 10 μL PBS were injected intradermally into mice anaesthetized with Ketamine using a 10μl Hamilton Syringe. In some experiments, MTB Erdman strain was used with same results. For aerosol infections, a frozen stock of Kanamycin-resistant Mtb H37Rv was diluted and used to infect mice in an aerosol infection chamber (Glas-Col). For high dose mEmerald infections, a deposition of 3000-5000 CFU was targeted. For Ultra low dose infections, a deposition of 1-3 CFU was targeted accepting a rate of approximately 30% uninfected animals. Bacterial load in the lungs was determined by plating serial dilutions from homogenized lungs. MTB was plated on Kanamycin-containing plates and antibiotic-free plates to distinguish the origin of the infection (intradermal versus aerosol infection).

### Heterologous challenges

For experiments with bacterial burden as the endpoint, 10^5^ Listeria monocytogenes were injected i.v. and CFU in the spleen measured 48 hours later. For experiments with an early response as endpoint, mice were infected with 10^7^ Listeria monocytogenes and sacrificed 2 hours after i.v. infection. The B16-F10 (ATCC® CRL-6475™) melanoma cell line was purchased from ATCC and expanded according to the supplier’s instructions. For the melanoma challenges, 10^5^ melanoma cells per mouse were injected i.v. After 10 days, mice were sacrificed, lungs were extracted and bleached in Fekete’s solution following a published protocol ^47^, and the metastases were counted by an investigator blinded to the identity of each mouse.

### Cell Isolation, Analysis, and Sorting

For i.v. labeling, PE labeled CD45.2 was injected i.v. 10 minutes prior to harvest. Single-cell suspensions of lung cells were prepared by Liberase Blendzyme 3 (Roche) digestion of perfused lungs as previously described^48^. Cells from spleens were prepared as previously described (Spleen digestion protocol, Miltenyi Biotec). Fc-receptors were blocked with anti-CD16/32 (clone 2.4G2). Cells were suspended in 1×PBS (pH 7.4) containing 2.5% FBS and stained at saturating conditions with antibodies against various epitopes (see Table S4). Samples were fixed in 2% (vol/vol) paraformaldehyde and analyzed using a LSRII flow cytometer (BD) and FlowJo software (Tree Star, Inc.). Previously published gating strategies were followed^49, 50^. In some experiments, alveolar macrophages were isolated from suspensions of lung cells using a BD Aria II cell sorter. Gating strategies are presented in Supplemental Figure 12 (T cells), Supplemental Figure 13 (Circulating monocytes), Supplemental Figure 14 (Spleen myeloid cells), and Supplemental Figure 15 (Lung myeloid cells).

### CD11b^+^ cell enrichment and ex vivo stimulation

Single cell suspensions prepared as described above from pooled lungs and spleens were positively enriched for CD11b^+^ using magnetic beads (Miltenyi Biotec). The cells were re-stimulated with TLR-agonists (LPS 10 ng/mL, PAM3 300 ng/mL, R848 100 ng/mL) and supernatants were collected 6 hours after re-stimulation and assayed by ELISA for TNF-α.

### Measurement of total protein

Cytokines were measured by the Cytokine & Chemokine 36-plex Mouse ProcartaPlex panel (with a Leptin assay added) and IFN alpha/IFN beta 2-Plex Mouse ProcartaPlex Panel (for a total of 39 analytes) using a Luminex Bio-Plex 200 analyzer per manufacturer instructions. Total protein levels were normalized using a commercial BCA assay from Thermo Scientific.

### Detection of Mtb-Specific T Cells

For direct detection of Mtb-specific cells, APC-labeled MHC class II tetramers (I-Ab) containing the immunodominant epitope of the ESAT-6 protein of Mtb (ESAT-64–17:I-Ab) were made in house. Pacific Blue-labeled MHC class I tetramers containing the stimulatory residues of the TB10.4 protein of Mtb (TB10.4 4–11:Kb) were obtained from the National Institutes of Health Tetramer Core Facility. Tetramer staining on single-cell preparations was carried out as described previously^51^.

### Immunohistochemistry

Tissue sections were formalin-fixed and paraffin-embedded. H&E Staining was carried out using standard protocols. An independent, board-certified pathologist reviewed and scored the tissue sections, blinded to their experimental group membership.

### RNAseq

RNA isolation was performed using TRIzol (Invitrogen), two sequential chloroform extractions, Glycoblue carrier (Thermo Fisher), isopropanol precipitation, and washes with 75% ethanol. RNA was quantified with the Bioanalyzer RNA 6000 Pico Kit (Agilent). cDNA libraries for alveolar macrophages were constructed and amplified using the SMARTer Stranded Total RNA-Seq Kit v2 – Pico Input Mammalian (Clontech) per the manufacturer’s instructions. cDNA for whole-blood was prepared using the TruSeq Stranded mRNA kit (Illumina, CA, USA). Libraries were amplified and then sequenced on an Illumina NextSeq (2 × 75, paired-end). Stranded paired-end reads of length 76 were preprocessed: For the Pico Input prep, the first three nucleotides of R2 (v2 kit) were removed as described in the SMARTer Stranded Total RNA-Seq Kit – Pico Input Mammalian User Manual (v2: 063017); Read ends consisting of 50 or more of the same nucleotide were removed. The remaining read pairs were aligned to the mouse genome (mm10) + Mtb H37Rv genome using the GSNAP aligner (v. 2016-08-24) allowing for novel splicing. Concordantly mapping read pairs (average 10-20 million / sample) that aligned uniquely were assigned to exons using the subRead (v. 1.4.6.p4) program and gene definitions from Ensembl Mus_Musculus GRCm38.78 coding and non-coding genes. Differential expression was calculated using the edgeR package from bioconductor.org. False discovery rate was computed with the Benjamini-Hochberg algorithm. Raw and processed data are deposited in GEO (GSE126355).

### ATACseq

The ATAC-seq protocol developed by with modifications for PFA fixed cells as described by Chen et al. (2016) was used^52^. Libraries were sequenced on a NextSeq 500 (Illumina) using a 150 (paired-end 2×76bp) cycle mid-output kit. Unique sequence read pairs were aligned to the combined *Mus musculus* (mm10) and *Mycobacterium tuberculosis* (H37Rv) genomes using the GSNAP aligner^53, 54^(v. 2016-08-24) after stripping off adapter sequences in a pairwise manner. Only pairs that aligned uniquely and concordantly to non-mitochondrial mouse chromosomes were retained. Start and end positions of the sequences were adjusted to extend 4 and 5 base pairs respectively to account for transposase adapter insertion (see Buenrostro et al. (2013)^55^). Peak calling was performed with MACS2 (2.1.0) ^56^using the start and end locations of the pairs to define fragment lengths. “Blacklisted” regions of known artificially high signal as defined by the ENCODE project were filtered out of peak regions (https://sites.google.com/site/anshulkundaje/projects/blacklists) ^57^. Bigwig files for each biological group were generated by running MACS2^56^ (https://github.com/taoliu/MACS) peak calling on combined alignments from all samples in the group and outputting a normalized bedgraph file followed by file conversion using the bedGraphToBigWig program (genome.ucsc.edu). The R package DiffBind^58^was used to define consensus peak regions across samples and assign counts. Differential peak counts and significance were computed using the R package edgeR^59, 60^. Raw and processed data are deposited in GEO (GSE126355).

### Whole blood transcriptome deconvolution analysis

Computational deconvolution of cell type proportion from whole-blood gene expression data was performed as previously described^61^. This approach used the assumption that whole-blood gene expression can be modelled as a mixture of gene abundances attributable to each individual cell type in the sample. Using a basis matrix containing a representative expression profile for each relevant cell type, cell proportions were estimated by fitting a model of the form:

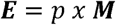

where **E** represents the whole-blood gene expression of a sample, **M**, the representative basis matrix, and *p*, the proportion of each cell type in the basis matrix, which will be estimable from the model coefficients when fit error is minimized. For human samples, the *immunoStates* matrix (*Valliana et al*) basis matrix was selected. This matrix has been explicitly designed to be transferrable across technology platforms, and has been successfully applied previously to deconvolute human whole-blood gene expression in an LTBI cohort^21^. For mouse samples, the *ImmuCC* basis matrix was used^22^.

Cell deconvolution was performed by fitting linear models (R: lm() function) to each Z-score normalized sample expression, as a function of the Z-score normalized basis matrix. Cell proportions were then directly estimated as the linear model coefficients, scaled to sum to 1. Mouse data are deposited in GEO (GSE126355); human data were obtained from^24^.

### Statistical Analysis

Significance was determined using an unpaired two-tailed Student’s *t*-test unless otherwise specified.

## Data Availability

Raw and processed RNA-seq and ATAC-seq data are deposited in GEO (GSE126355) (Figure 1A, Figure 4, Supplemental Figure 2, Supplemental Figure 10). Data for Figure 1E and Supplemental Figure 2 was obtained from GEO (GSE19491, GSE19444, GSE19443, GSE19442, GSE19439, GSE19435).

## ACKNOWLEDGMENTS

We thank the staff from Center for Infectious Disease Research for animal care; and Dr. Piper Treuting and Brian Johnson, University of Washington Immunohistochemistry core, for histological services. We acknowledge the NIH Tetramer Core Facility (Contract HHSN272201300006C) for provision of tetramers. We thank Martina Sanlorenzo and Igor Vujic for help with the melanoma model. Research reported in this publication was supported by the National Institute of Allergy and Infectious Diseases (Grant numbers U19AI135976, U19AI100627, and R01AI032972). JN was supported by Swiss National Foundation (SNF) grant #P300PB_164742.

## Supplemental figure legends

**Supplementary Figure 1: Summary of infected tissues in the LTBI model** Mice were inoculated intradermally in the ear and bacterial burden in the superficial cervical lymph nodes measured 10 days, 42 days, and 1 year by CFU assay. Plots depict the fraction of mice with detectable CFUs of Mtb in spleen across all timepoints (4-5 mice/timepoint).

**Supplementary Figure 2: Deconvolution analysis of whole-blood RNA-seq measurements in mice and humans** *Left panel:* Deconvolution analysis of whole-blood RNA-seq measurements to estimate changes in the proportions of circulating immune cell subsets 42 days following establishment of LTBI. (See Methods) (4 mice/condition) *Right panel:* Deconvolution analysis of whole-blood RNA-seq measurements on 69 individuals with LTBI and 117 uninfected individuals. Error bars represent the SD. Significance was determined by *t*-Test with the Benjamini and Hochberg correction for multiple testing (FDR). * FDR < 0.05.

**Supplementary Figure 3: Proportions of M1 and M2 macrophages in humans** Proportions of M1 and M2 macrophages in uninfected individuals (n=117, grey) and individuals with LTBI (n=69, blue) as determined by deconvolution analysis of whole blood RNA-seq measurements.

**Supplementary Figure 4: Protective effect of LTBI assessed at 6 weeks following aerosol challenge.** LTBI was established as described in the main text and mice challenged with 100-200 CFU of Mtb H37Rv via aerosol after 10-14 days (“Early”, 2 replicates) or after 8-10 weeks (“Late”, 4 replicates). Bacterial burden in the lung was measured by CFU assay. LTBI mice had on average 18.4-fold (CI: 10.6-26.3) fewer bacteria in the lung as compared to controls. All experiments were significant with *p* < 0.05. *t*-Tests was used for testing significance. Error bars represent SEM.

**Supplementary Figure 5: Protective efficacy of BCG Pasteur** Mice were immunized intradermally with 1×10^6^ CFU BCG Pasteur and challenged with 100 CFU Mtb H37Rv after 2 months. Bacterial burden in the lung was measured by CFU assay at days 10, 42, and 100 following aerosol challenge (n = 4-5 mice/group/time-point).

**Supplementary Figure 6: Correlation between cytokines levels in lungs and spleens of LTBI mice** LTBI was established as described in the main text and the abundances of selected cytokines and chemokines in the lungs and spleen measured by multiplexed immunoassay at day 42 following inoculation. Plot depicts the levels of analytes detected in both tissues. Each point represents the average of 5 biological replicates.

**Supplementary Figure 7: Proportions of immune cells in the lungs of naive and LTBI mice.** LTBI was established as described in the main text. At 8 weeks the absolute number of CD45^+^ cells in whole-lung homogenates was measured by flow cytometry using counting beads (left panel). The relative proportions of various immune cell populations were measured by flow cytometry (right panel). Representative data are shown for 1 of 2 independent experiments with 4-5 mice/condition. *t*-Tests was used for testing significance. Error bars represent SD.

**Supplementary Figure 8: Distribution of Mtb infection across pulmonary immune cells.** Flow panels show mEmerald+ MTB infected AMs gated on CD11c^+^CD64^+^SiglecF^+^CD45^+^CD11b^int^ cells 24h after high dose infection. LTBI and control mice were infected with ∼2000-4000 CFU of mEmerald-expressing Mtb via aerosol and infected cell types in BAL fluid were assessed at 24 hours by flow cytometry. Error bars represent SD.

**Supplementary Figure 9: Bacterial burden in the lung following high-dose aerosol challenge.** Control and LTBI mice were infected with ∼2000-4000 CFU of Mtb and bacterial burden in the lung measured by CFU assay 10 days following challenge. Representative data from 1 of 2 independent experiments with 4-5 mice/condition. Significance was assessed by Student’s *t*-Test. Error bars represent the mean with SD; * *p* < 0.05.

**Supplementary Figure 10: ATAC-seq on AM from naïve/latent mice at baseline.** Alveolar macrophages from LTBI and control mice (n=3/condition) were isolated from BAL fluid by FACS and ATACseq was performed following a published protocol^52^. Plot depicts FDR vs. difference in chromatin accessibility between control and LTBI for 45,458 genomic regions (See Methods). Red dots repreent promotor regions (See Methods).

**Supplementary Figure 11: T cells from treated LTBI mice fail to expand upon infection.** The fraction of CD3^+^CD8^+^TB10.4^+^ T cells in whole lung homogenates prior to challenge and at 10 days following challenge was determined by flow cytometry. Significance was assessed by Student’s *t*-Test. Error bars represent the mean with SD; * *p* < 0.05.

## Contributions

JN, AD, EG, KU and AA designed the experiments. JN, AJ, DM, GSO, AR, CP, JD did the animal work. AD, EG and FD analyzed the RNAseq data and performed the computational analysis. SS performed the ATAQseq experiment. JN, GSO, AD and EG wrote the manuscript. All authors reviewed and/or provided feedback on the manuscript.

