## Supplemental_figures for "Latent *Mycobacterium tuberculosis* infection provides protection for the host by changing the activation state of the innate immune system"

### Supplemental Figure 1

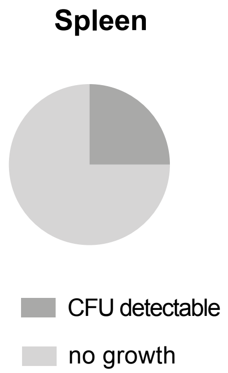


### Supplemental Figure 2

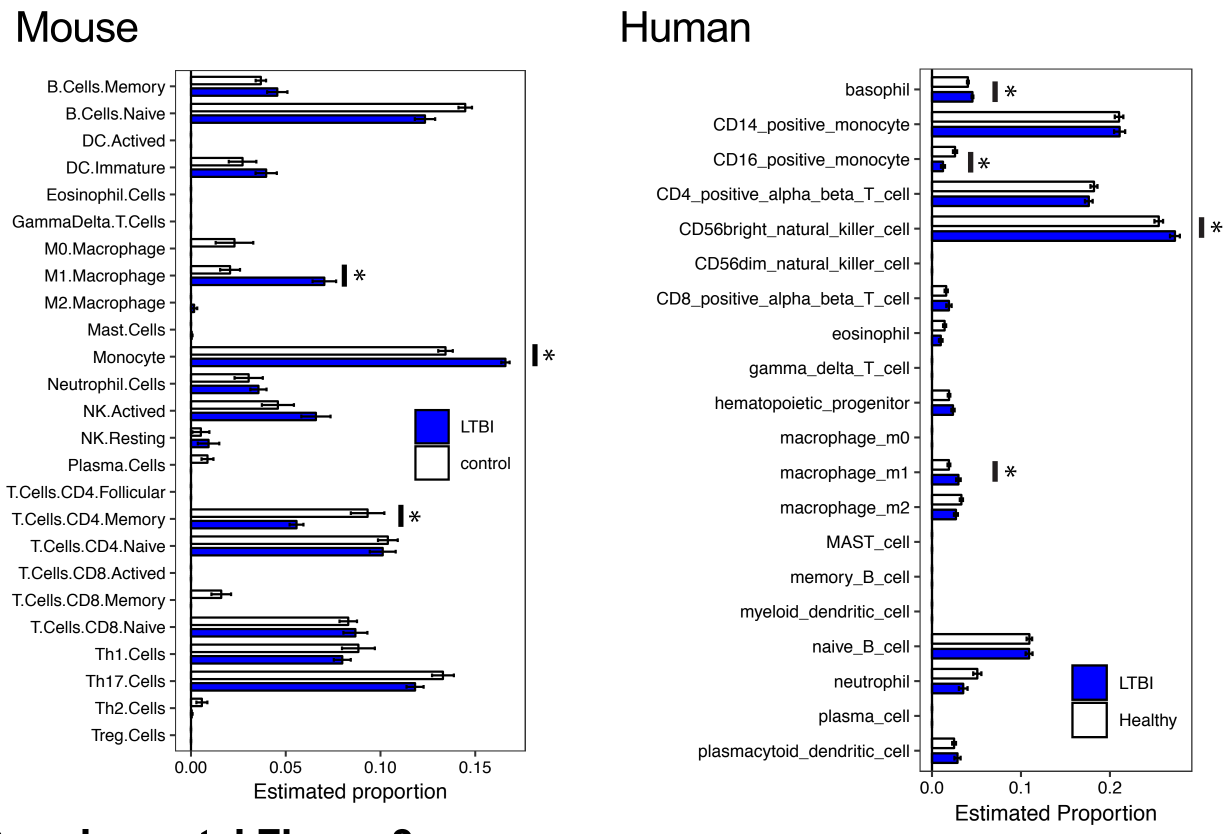


### Supplemental Figure 3

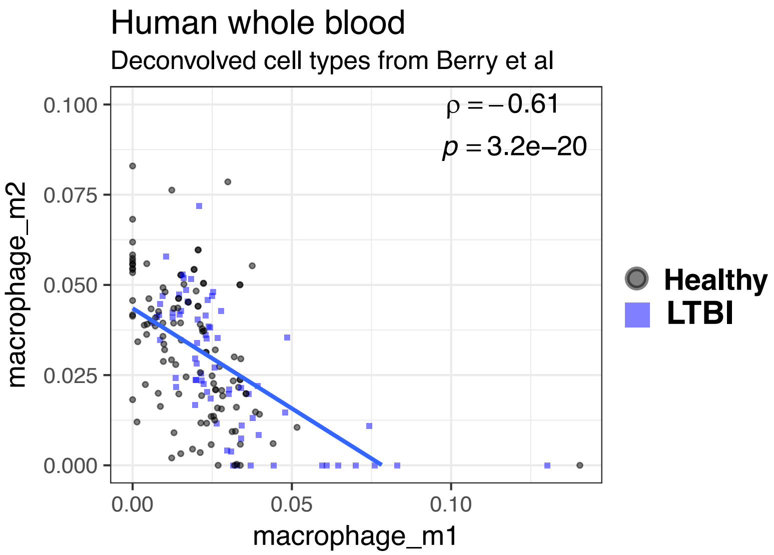

#### Supplemental Figure 4

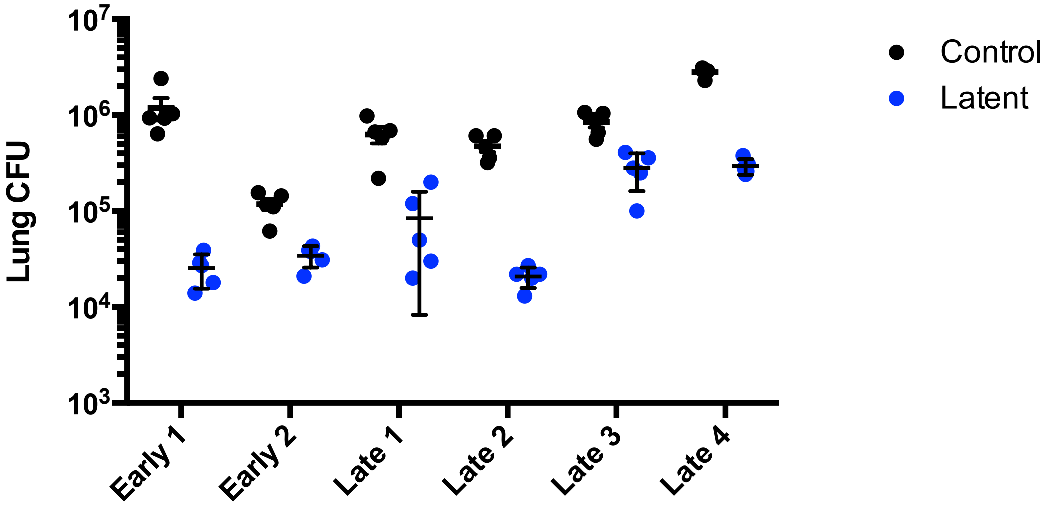


#### Supplemental Figure 5

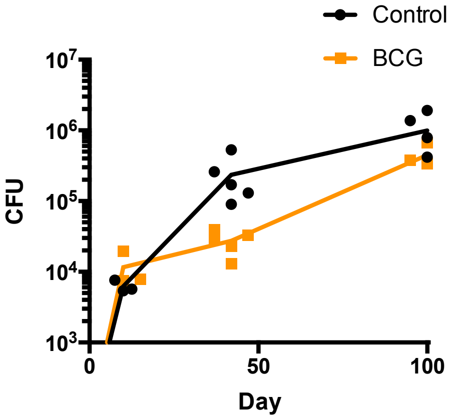

#### Supplemental Figure 6

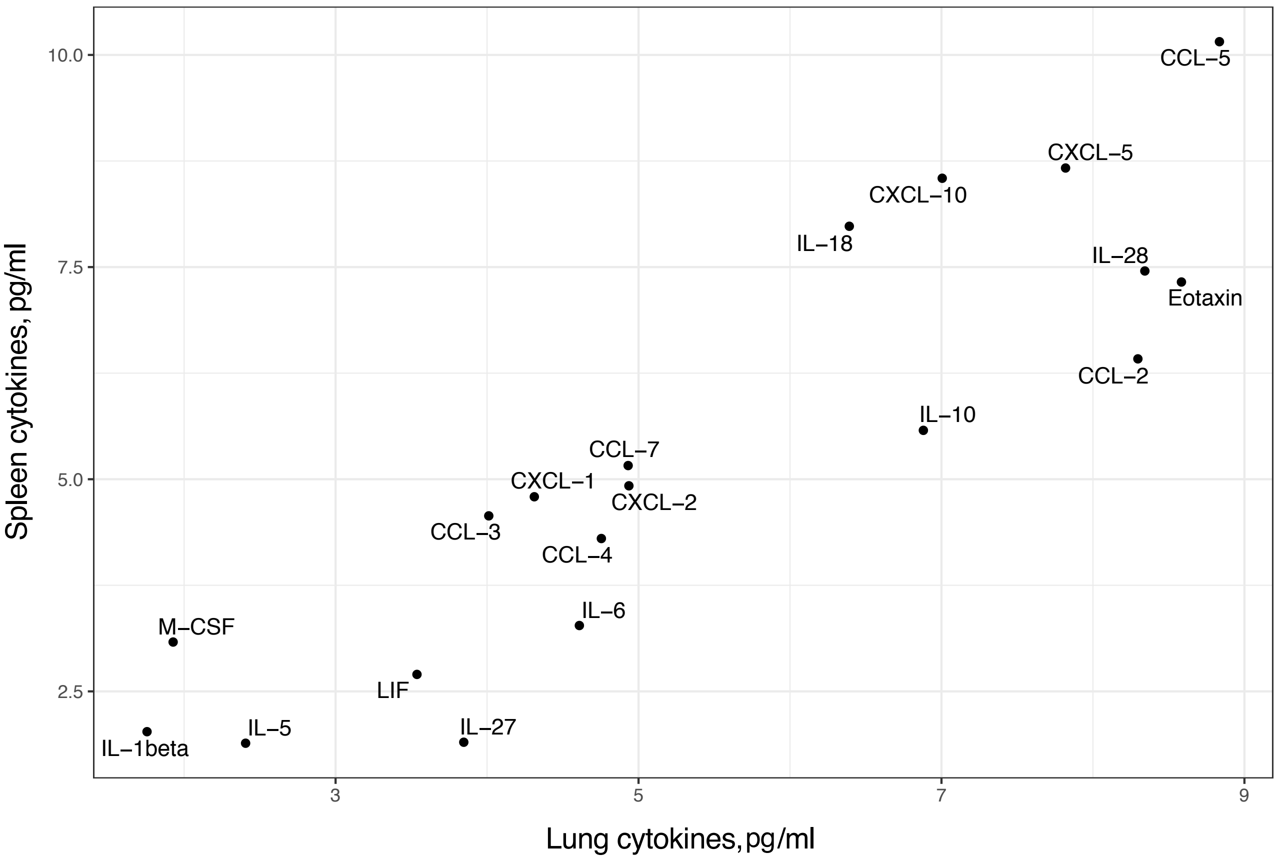


#### Supplemental Figure 7

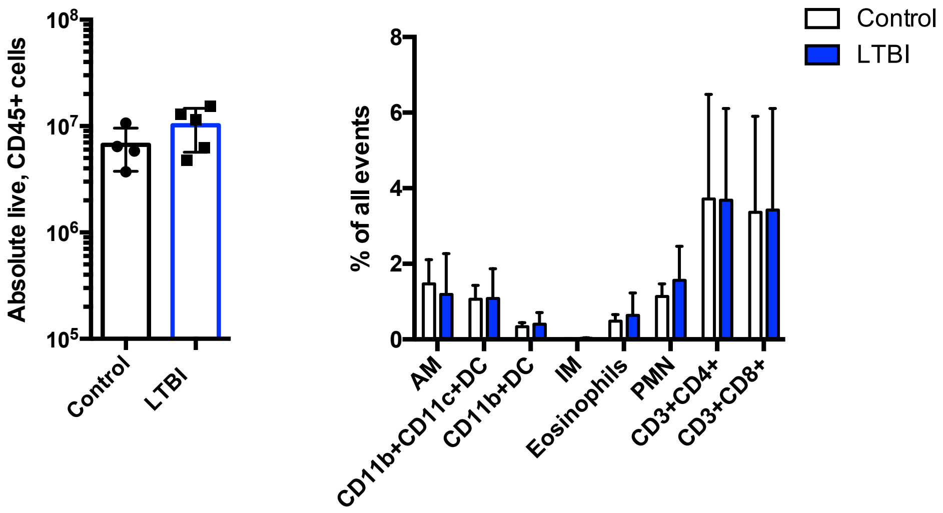

#### Supplemental Figure 8

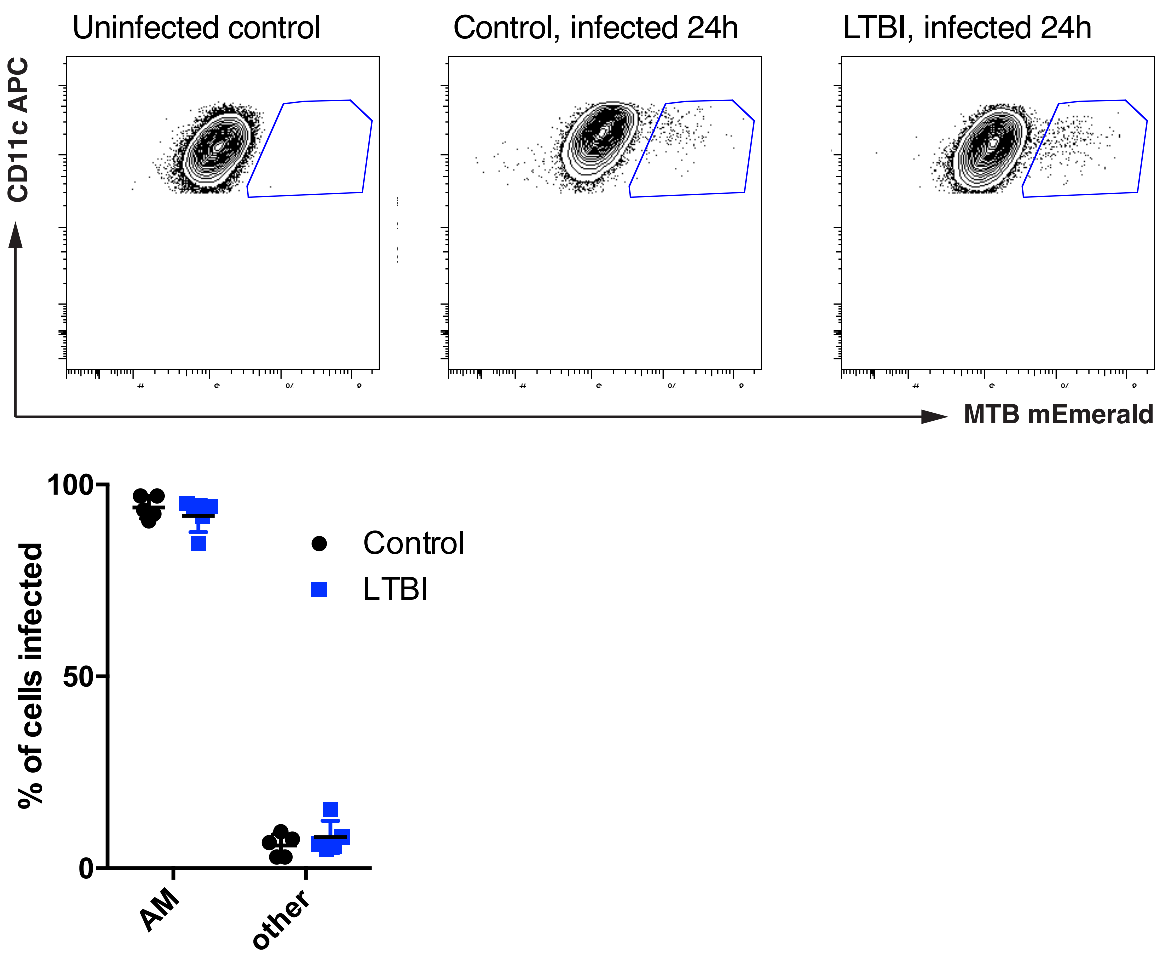


#### Supplemental Figure 9

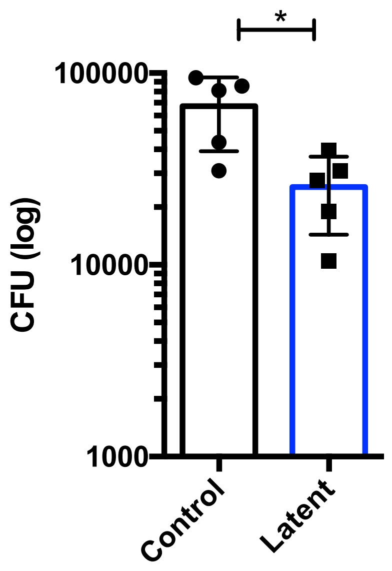


#### Supplemental Figure 10

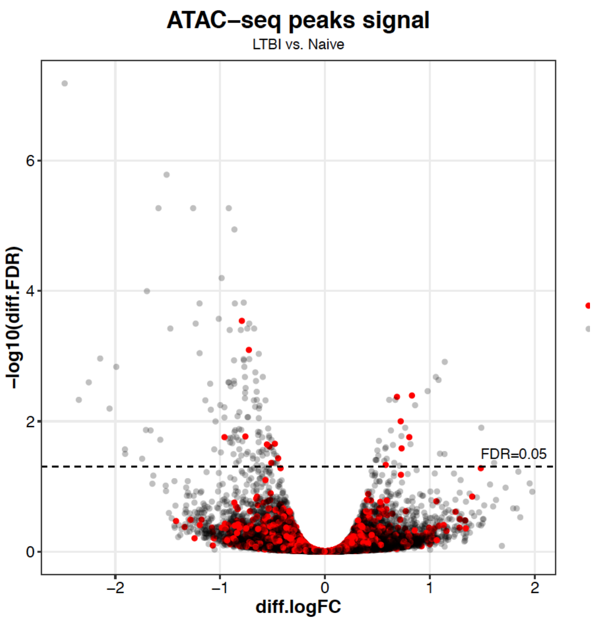

#### Supplemental Figure 11

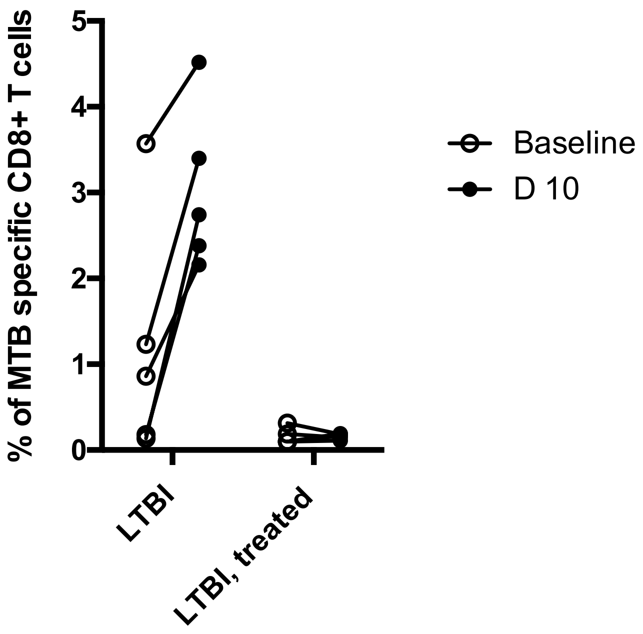

#### Supplemental Figure 12: T cell gating strategy

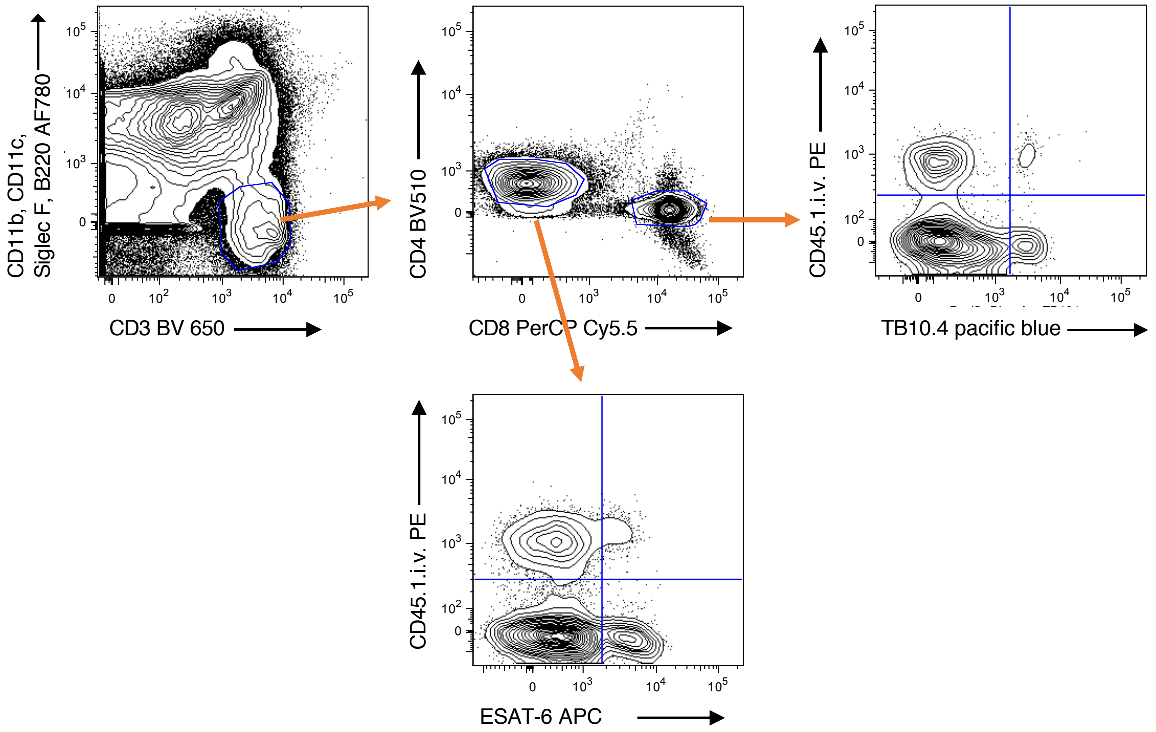


#### Supplemental Figure 13: Monocyte from peripheral blood gating strategy

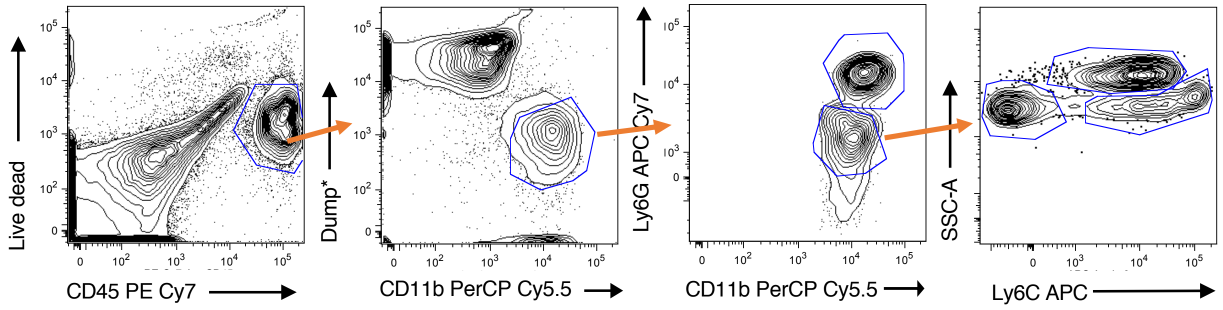


*B220, TCRab, NK1.1., CD11c PE

#### Supplemental Figure 14: Spleen myeloid panel gating strategy

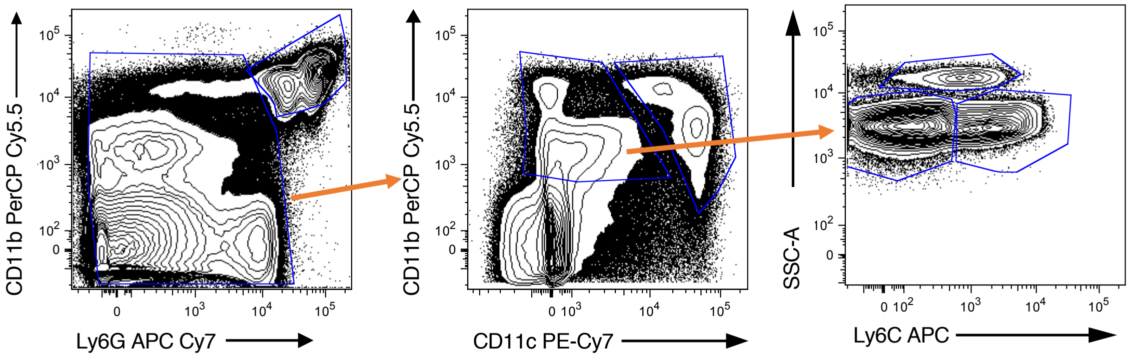


#### Supplemental Figure 15: Lung myeloid panel gating strategy

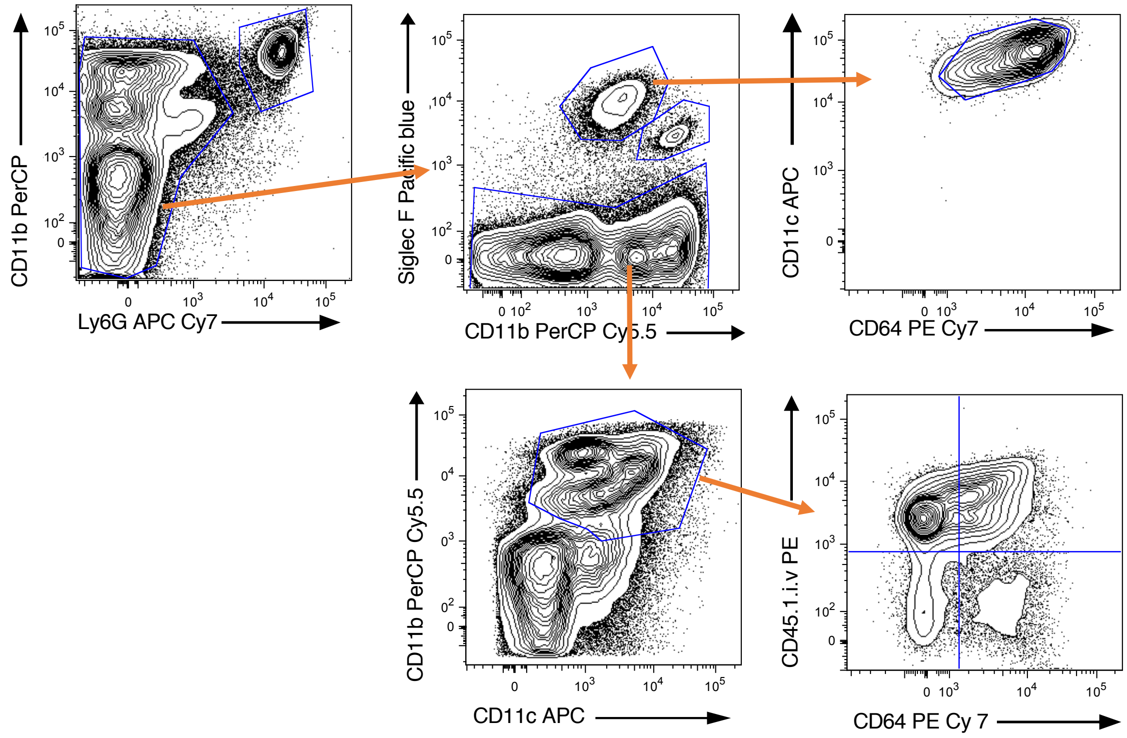
